# Bonsai: An event-based framework for processing and controlling data streams

**DOI:** 10.1101/006791

**Authors:** Gonçalo Lopes, Niccolò Bonacchi, João Frazão, Joana P. Neto, Bassam V. Atallah, Sofia Soares, Luís Moreira, Sara Matias, Pavel M. Itskov, Patrícia A. Correia, Roberto E. Medina, Lorenza Calcaterra, Elena Dreosti, Joseph J. Paton, Adam R. Kampff

## Abstract

The design of modern scientific experiments requires the control and monitoring of many parallel data streams. However, the serial execution of programming instructions in a computer makes it a challenge to develop software that can deal with the asynchronous, parallel nature of scientific data. Here we present Bonsai, a modular, high-performance, open-source visual programming framework for the acquisition and online processing of data streams. We describe Bonsai’s core principles and architecture and demonstrate how it allows for flexible and rapid prototyping of integrated experimental designs in neuroscience. We specifically highlight different possible applications which require the combination of many different hardware and software components, i0ncluding behaviour video tracking, electrophysiology and closed-loop control of stimulation parameters.

## Introduction

Modern scientific experiments crucially depend on the control and monitoring of many parallel streams of data. Multiple measurement devices ranging from video cameras, microphones and pressure sensors to neural electrodes, must simultaneously send their data in real-time to a recording system. General purpose digital computers have gradually replaced many of the specialized analog and digital technologies used for this kind of data acquisition and experiment control, largely due to the flexibility of programming and the exponential growth in computing power. However, the serial nature of programming instructions and shared memory makes it a challenge, even for experienced programmers, to develop software that can elegantly deal with the asynchronous, parallel nature of scientific data.

Another challenge arises from the need for software integration. Each hardware vendor provides their own set of drivers and programming interfaces for configuring and acquiring data from their devices. In addition, the growth of the open-source software movement has greatly increased the number of freely available technologies for different data processing domains. Integration of all these diverse software and hardware components remains a major challenge for researchers.

These difficulties lead to increased development times in setting up an experiment. Moreover, it requires the experimenter to undergo specialized training outside his domain of research. This limits the ability to rapidly prototype and try out new designs and can quickly become a constraining factor in the kinds of questions that are amenable to scientific investigation.

Here we describe Bonsai, an open-source visual programming framework for processing data streams. The main goal of Bonsai is to simplify and accelerate the development of software for acquisition and processing of the many heterogeneous data sources commonly used in (neuro)scientific research. We expect to facilitate the fast implementation of state-of-the-art experimental designs in many labs and also allow easy exploration of new designs. The framework has already been successfully used in many applications. In this work we emphasize its use in neuroscience for monitoring and controlling behaviour and physiology experiments.

## Results

### 1. Architecture

Bonsai was developed on top of the Reactive Extensions for the .NET framework (Rx)^1^. Rx represents asynchronous data streams using the notion of an observable sequence. As the name implies, elements in an observable sequence follow one after the other. The name observable simply specifies that the way we access elements in the data stream is by listening to (i.e. observing) the data as it arrives, in contrast with the static database model, in which the desired data is enumerated. Rx also provides many built-in operators that transparently handle the parallelism arising from the combination of multiple data sources, making it an ideal basis on which to build frameworks for reactive and event-driven software.

In Bonsai, observable sequences are created and manipulated graphically using a dataflow^2,3^ representation (Fig. 1a). Each node in the dataflow represents an observable sequence. Nodes can be classified as either observable sources of data or combinators. Sources provide access to raw data streams, such as images from a video camera or signal waveforms from a microphone or electrophysiology amplifier. Combinators represent observable operators that bring together one or more sequences, and can be further specialized into transforms and sinks depending on how they manipulate their inputs. Transforms change the incoming data elements from a single input sequence. An example would be taking a sequence of numbers and producing another sequence of numbers containing the original elements multiplied by two. Sinks, on the other hand, simply introduce processing side-effects without modifying the original sequence at all. One example would be printing each number in the sequence to a text file. The act of printing in itself changes nothing about the sequence, which continues to output every number, but the side-effect will generate a useful action. If a combinator is neither a transform nor a sink, it is simply called combinator. The *Sample* combinator illustrated in (Fig. 1a) takes two data sequences and produces a new sequence where elements are sampled from the first sequence whenever the second sequence produces a new value. In this example, we use *Sample* to extract and save single images from a video stream whenever a key is pressed.

**Figure 1:**
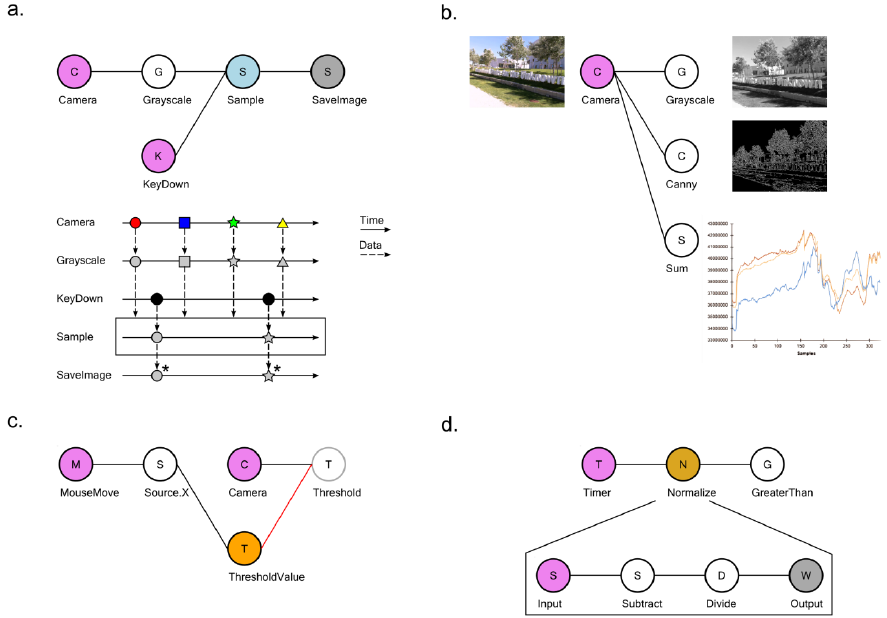
Examples of dataflow processing pipelines using Bonsai. **a.** Taking grayscale snapshots from a camera whenever a key is pressed. Top: graphical representation of the Bonsai dataflow for camera and keyboard processing. Data sources are coloured in violet; transform operators in white; combinators in light blue. Bottom: marble diagram showing an example execution of the dataflow. Coloured tokens represent frames arriving from the camera. Black circles represent key press events from the keyboard. Asterisks indicate saving of images to permanent storage. **b.** Visualizing image processing filters. The inset next to each node represents the corresponding Bonsai data visualizer. **c.** Dynamic modulation of an image processing threshold using the mouse. The x-coordinate of mouse movements is used to directly set the externalized ThresholdValue property (orange). The updated threshold value will be used to process any new incoming images. **d.** Grouping a set of complex transformations into a single node. In the nested dataflow, the source represents incoming connections to the group and the sink represents the group output.

A common requirement when designing and manipulating dataflows is the ability to visualize the state of the data at different stages of processing. We have included a set of visualizers to assist debugging and inspection of data elements, including images and signal waveforms (Fig. 1b). These visualizers are automatically associated with the output data type of each node and can be launched at any time in parallel with the execution of the dataflow. Furthermore, it is often desirable to be able to manipulate processing parameters online for calibration purposes. Each node has a set of properties which parameterize the operation of that particular source or combinator. This allows, for example, changing the cutoff frequency of a signal processing filter, or setting the name of the output file in the case of data recording sinks. We have also included the possibility of externalizing node properties into the dataflow (Fig. 1c). Externalizing a property means pulling out one of the parameters into its own node in the dataflow, making it possible to connect the output of another node to the exposed property. This allows for the dynamic control of node parameters.

Finally, we have built into Bonsai the ability to group nodes hierarchically. In its simplest form, this feature can be used to encapsulate a set of operations into a single node which can be reused elsewhere (Fig. 1d). This is similar to defining a function in a programming language and is one of the ways to create new reactive operators in Bonsai. Any named externalized properties placed inside an encapsulated dataflow will also show up as properties of the group node itself. This allows for the parameterization of nested dataflows and increases their reuse possibilities. In addition, encapsulated dataflows are used to specify more complicated, yet powerful, operators such as iteration constructs that allow for the compact description of complex data processing scenarios, but that can be cumbersome to specify in pure dataflow visual languages^3^ (see below).

Bonsai was designed to be a modular framework, which means it is possible to extend its functionality by installing additional packages containing sources and combinators developed for specific purposes. New packages can be written by using C# or any of the .NET programming languages. Python scripts (via IronPython^4^) can be embedded in the dataflow as transforms and sinks, allowing for rapid integration of custom code. All functionality included in Bonsai was designed using these modular principles, and we hope to encourage other researchers to contribute their own packages and thereby extend the framework to other application domains. At present, the available packages include computer vision and signal processing modules based on the OpenCV^5^ library. Drivers for several cameras and interfaces to other imaging and signal acquisition hardware were integrated as Bonsai sources and sinks, including support for Arduino^6^ microcontrollers, serial port devices and basic networking using the OSC^7^ protocol. Given the specific applications in the domain of neuroscience, we also integrated a number of neuroscience technology packages. The Ephys package, for example, builds on the Open Ephys initiative for the sharing of electrophysiology acquisition hardware^8^ by providing support for the Rhythm open-source USB/FPGA interface (Intan Technologies, US). This means that the next generation tools for electrophysiology can be used inside Bonsai today and the acquired data integrated with many other available data streams into a powerful, flexible and standardized experimental neuroscience platform.

### Advanced Operators

The most common and simple application of Bonsai is the acquisition and processing of simple, independent data streams. However, for many modern experiments, simple acquisition and storage of the data is often not sufficient. For example, it can be convenient to only record data aligned on events of interest, such as the onset of specific stimuli. Furthermore, neuroscience experiments often progress through several stages, especially for behavioural assays, where controlled conditions need to vary systematically across different sessions or trials. In order to enforce these conditions, experiments need to keep track of which stage is active and use that information to update the state of control variables and sensory processing. These requirements often cannot be described by a simple linear pipeline of data, and require custom code to handle the complicated logic and bookkeeping of experimental states. Below we describe a set of advanced Bonsai operators that can be used to flexibly reconfigure data processing logic to cover a larger number of scenarios. These operators and their applications are all built on the single idea of slicing a data stream into sub-sequences, called windows, which are then processed independently and potentially in parallel (Fig. 2).

**Figure 2:**
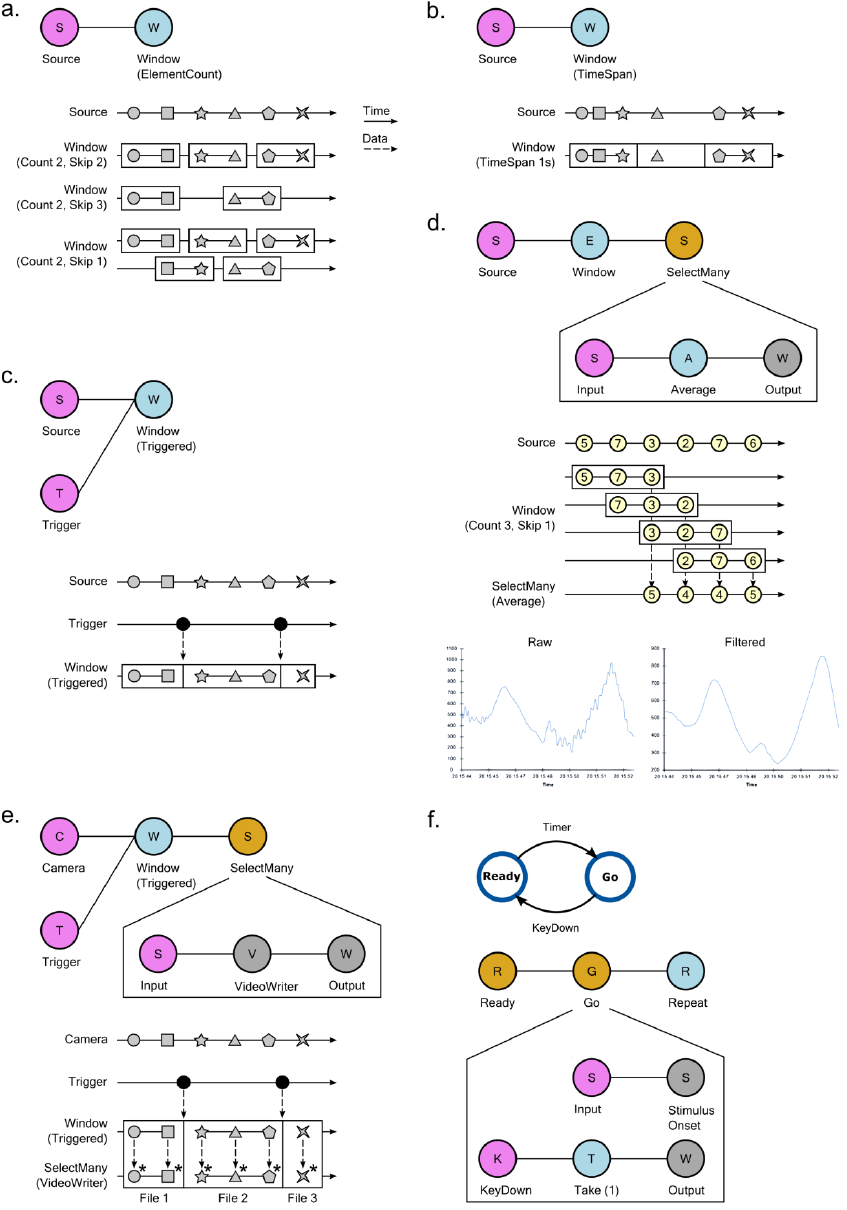
Using slicing and window processing combinators in Bonsai. **a.** Creating data windows using element count information. The Count and Skip parameters specify the size of each window and the number of elements to skip before creating a new window, respectively. Top: graphical representation of the Bonsai dataflow used for slicing. Bottom: marble diagram showing the behaviour of the operator for different values of the parameters. The boundaries of each window are indicated by the enclosing rectangles. **b.** Creating data windows using timing information. Time is split into intervals of equal fixed duration indicated by the enclosing rectangles. Each interval defines a window and data elements are assigned to each window based on the interval that is active at the time of their arrival. Top: Bonsai dataflow. Bottom: marble diagram. **c.** Creating data windows using an external trigger. The boundaries of the created windows are defined by the timing of events produced by the trigger source. Top: Bonsai dataflow. Bottom: marble diagram. **d.** Moving average of a signal source using windows. Sliding windows of the data are created based on element count information. Top: Bonsai dataflow. The dataflow encapsulated in *SelectMany* specifies the processing done on each window. In this case, the average value of each window sequence is computed. Middle: marble diagram. As soon as each window is completed, its average value is merged into the result sequence. Bottom: example signal trace before and after the filtering. **e.** Online splitting of video recordings into different files based on an external trigger. Top: Bonsai dataflow. Notice that the *VideoWriter* sink is included inside the *SelectMany* combinator. Bottom: marble diagram. At the start of each window, a new movie file is created. Frames in each window are encoded in the corresponding file. **f.** Implementing state-machines using Bonsai window operators. Top: state-machine schematic of a task designed to measure response times. In the Ready state, the stimulus is off. When entering the Go state, the stimulus is turned on. At the end of each trial, the system goes back to the initial state. Bottom: graphical representation of the equivalent Bonsai dataflow. The *SelectMany* combinator is used to specify the behaviour and transitions of each state. The *Take* combinator truncates a sequence to include only a specified number of initial elements. In this case, only the first element is included. The *Repeat* combinator restarts a sequence when no more elements are produced (see text).

Bonsai provides different combinators that allow the creation of these sub-sequences from any observable data stream, using element count information, timing, or external triggers (Fig. 2a)–(Fig. 2c). The specific set of operations to apply on each window is described by encapsulating a dataflow inside a *SelectMany* group, as detailed in the signal processing example of Fig. 2d. The input source in this group represents each of the window sub-sequences, i.e. it is as if each of the windows is a new data source, containing only the elements that are a part of that window. These elements will be processed as soon as they are available by the encapsulated dataflow. Windows can have overlapping common elements, in which case their processing will happen concurrently. The processing outputs from each window are merged together to produce the final result. In the case of Fig. 2d, past and future samples are grouped in windows to compute a moving average of the signal through time, necessarily time-shifted by the number of future samples that are considered in the average.

The processing of the elements of each window happens independently, as if there was a new isolated dataflow running for each of the sequences. We can exploit this independence in order to dynamically turn dataflows on and off during an experiment. In the video splitting example of Fig. 2e, we use an external trigger source to chop a continuous video stream into many small video sequences, aligned when the trigger fired. We then nest a *VideoWriter* sink into the *SelectMany* group. The *VideoWriter* sink is used to encode video frames into a continuous movie file. It starts by creating the video file upon arrival of the first frame, and then encoding every frame in the sequence as they arrive. When the data stream is completed, the file is closed. By nesting the *VideoWriter* inside the *SelectMany* group, what we have effectively done is to create a new video file *for each* of the created windows. Whenever a new trigger arrives, a new clip is created and saving proceeds transparently to that video file.

More generally, we can use this idea to implement discrete transitions between different processing modes, and chain these states together to design complex control structures such as finite state machines (FSM). FSMs are widely used to model environments and behavioural assays in systems and cognitive neuroscience. One example is illustrated in Fig. 2f, where we depict the control scheme of a stimulus-response apparatus for a simple reaction time task. In this task, there are only two states: Ready and Go. In the Ready state, no stimulus is presented and a timer is armed. Whenever the timer fires, the task transitions into the Go state, and a stimulus is presented. The subject is instructed to press a key as fast as possible upon presentation of the stimulus. As soon as the key is pressed, the system goes back to the Ready state to start another trial. In a FSM, nodes represent states, e.g. stimulus availability or reward delivery, and edges represent transitions between states that are caused by events in the assay, e.g. a key press. In each state, a number of output variables and control parameters are set (e.g. turning on a light) which represent the behaviour of the machine in that state.

In the Bonsai dataflow model, dataflows encapsulated in a *SelectMany* group can be used to represent states in a FSM (Fig. 2f, bottom). Specifically, a state is activated whenever it receives an input event, i.e. the dataflow nested inside the state will be turned on. The dynamics of the nested dataflow determine the dynamics of the state. In the Go state presented in Fig. 2f, the activation event is used to trigger stimulus onset. In parallel, we start listening for the key press which will terminate the state. Conversely, for the Ready state we would trigger stimulus offset and arm the timer for presenting the next stimulus. An important difference between Bonsai dataflows and pure state machine models is that we do not allow for cycles in the dataflow. However, by taking advantage of the *Repeat* combinator we can restart a dataflow once it’s completed, allowing us to reset the state machine for the next trial.

Many of the control tasks in experiments have this sequential trial-based structure, which has allowed us to rapidly prototype complex behaviour assays such as closed-loop rodent decision making tasks simply by leveraging the flexibility of the data stream slicing operators.

### Applications

The validation of Bonsai was performed by using the framework to implement a number of application use cases in the domain of neuroscience (Fig. 3). The breadth of technologies at use in this field demands that modern experiments be able to handle many heterogeneous sources of data. Experimenters need to routinely record video and sensor data monitoring the behaviour of an animal simultaneously with electrophysiology, fluorescent reporters of neural activity or other physiological measures. Online manipulation and visualization of data is a fundamental part of the experiment protocol for many of the reported techniques. In the following we highlight some of these practical applications of Bonsai in more detail in order to illustrate both best practices and implementation challenges.

**Figure 3:**
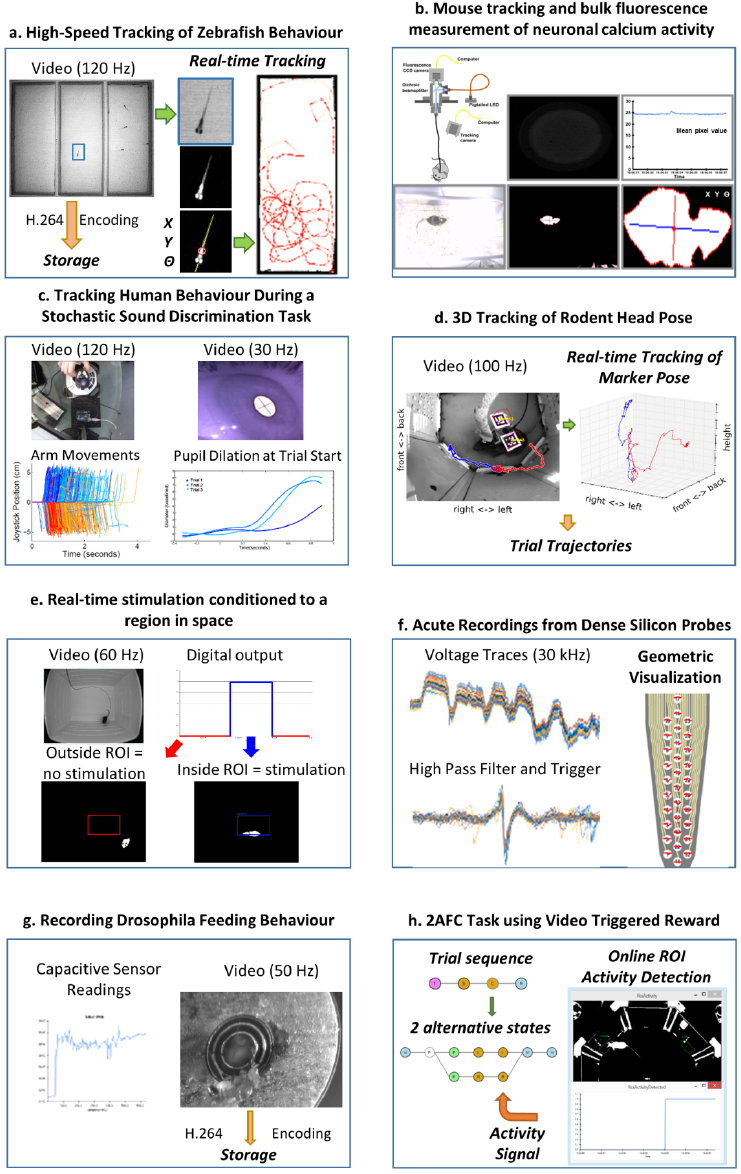
Example use cases of neuroscience experimental setups using Bonsai. **a.** High-speed tracking of zebrafish behaviour. Insets depict the image processing steps for segmenting the shape of a fish from the background and extracting its spatial location and orientation. Right: example trajectories extracted from an individual fish. **b.** Mouse tracking and bulk fluorescence measurement of neuronal calcium activity. Top insets: schematic of the fibre optic imaging setup for freely moving rodents with example fluorescence data frame and extracted fluorescence signal traces. Bottom insets: image processing steps for behaviour tracking of a mouse as the largest dark object in the video. **c.** Tracking human behaviour during a stochastic sound discrimination task. Left insets: arm movements on the joystick on each trial tracked by brightness segmentation of a bright LED. Right insets: extraction of pupil dilation by computing the length of the major axis of the largest dark object. **d.** 3D tracking of rodent head pose. Left inset: example video frame of a mouse carrying fiducial markers. A cube was rendered and superimposed on the image to demonstrate correct registration. Coloured traces show representative single trial trajectories of an individual marker, aligned on center poke onset. Red and blue refer to left and right choice trials, respectively. Right inset: Threedimensional plot of the same trajectories using isometric projection. **e.** Real-time stimulation conditioned to a region in space. Top insets: example raw movie frame and stimulation state. Red and blue indicate no stimulation and stimulation regimes, respectively. Bottom insets: example video frames where the mouse is either outside or inside the region of interest. **f.** Acute recordings from dense silicon probes. Left insets: example traces from raw amplified voltage signals and high-pass filtered spike triggered waveforms. Right inset: visualization of spike waveforms triggered on a single channel superimposed on the actual probe geometry. **g.** Recording drosophila feeding behaviour. Left inset: example trace of a single-channel capacitive signal from the flyPAD. Right inset: simultaneously recorded video of the fly feeding behaviour. **h.** 2AFC task using video triggered reward. Left inset: schematic of the reactive state machine used for controlling the task. Each state is represented by a nested dataflow. Branches represent possible transitions. Right inset: example thresholded activity from a single region of interest activated by the mouse.

One of the first main use cases driving the development of Bonsai was the automated online tracking of animal behaviour using video. The most common tracking application involves chaining together operators for image segmentation and binary region analysis to allow the extraction of the spatial location of an animal over time (Figs. 3a, 3b). The same technique can easily carry over to track different kinds of objects such as eyes or experimental manipulanda in human psychophysics experiments (Fig. 3c) provided adequate illumination contrast and the appropriate choice of a method for segmentation. These image processing tools can also be used to acquire and process physiological data in neural imaging setups, where it is now possible to record bioluminescent or fluorescent reporters of neural activity during behaviour. For example, Fig. 3b demonstrates simultaneous measurement of animal behaviour and neural activity using bulk fluorescence calcium imaging in the mouse brain recorded with a CCD sensor and a fiberoptic setup^9^.

Raw video data from modern high-resolution, high-speed cameras can be expensive and cumbersome to store. Online video compression and storage sinks were implemented taking advantage of parallelism to avoid frame loss. Video compression is processing intensive and can compromise data acquisition if reading the next frame has to wait for the previous frame to be fully encoded. One solution is to buffer incoming frames and compress them in parallel with the rest of the processing. By encapsulating this behaviour into a Bonsai sink, it became easy to incorporate video recording and compression functionality into any image processing pipeline (Figs. 3a–e, 3g-h).

While simple image processing techniques can easily extract continuous two-dimensional measures of animal location over time, it often becomes the case that the experimenter is concerned with tracking the detailed behaviour of specific features in the animal’s body, such as head pose. This is an essential component in neurophysiology or stimulation experiments in freely moving animals, where ongoing behaviour is the central constraint in interpreting neural responses and manipulations. However, identifying such features and reconstructing their position and orientation in 3D space is a challenging computer vision problem. A common solution is to use planar fiducial markers of known geometry^10^,^11^ (Fig. 3d). The computer vision research community has developed some open-source software solutions to this problem^10^, which have been integrated into Bonsai to allow the possibility of easily and flexibly incorporating online 3D fiducial tracking in video streams. This approach has been used successfully to record 3D head movements of a mouse under optogenetic stimulation in a decision-making task (Fig. 3d).

One final, but important application of video stream processing is in the development of closed-loop interfaces, where the actions of an animal directly modulate manipulations under the experimenter’s control. This kind of experiment requires fast online analysis of behaviour and physiological variables of interest that are subsequently coupled to hardware control interfaces. In Fig. 3e, real-time stimulation conditioned to a region in space was implemented by analysing the position of an animal in a square arena. Whenever the animal found itself inside a specified region of interest, a signal was sent to an Arduino controller which was then used to drive optogenetic stimulation of specific neural circuits.

Another key data type that is commonly processed by Bonsai dataflows is buffered time-series data. This type of data usually arises from audio, electrophysiology or other digital acquisition systems where multiple data samples, from one or more channels, are synchronously acquired, buffered and streamed to the computer. These buffers are often represented as data matrices, where rows are channels and columns represent individual data samples through time, or vice-versa. Support for simple band-pass filters, thresholding and triggering allowed us to build flexible spike detection and waveform extraction systems (Fig. 3f). Using Intan’s Rhythm API, we integrated into Bonsai support for a variety of next-generation electrophysiology devices using Intan’s digital amplifier technology, such as the Open Ephys acquisition system^8^ or Intan’s evaluation board (RHD2000, Intan Technologies, US). This system was successfully used to acquire and visualize simultaneous recordings from dense silicon probes where spikes from a loose-patch juxtacellular pipette were used as triggers to align and extract waveform data appearing on the multi-channel extracellular probe. Responses from every silicon probe site could then be superimposed on an accurate rendition of the probe geometry in real-time.

The ability to rapidly integrate new modules allowed us to support the development and crossvalidation of new tools for behavioural neuroscience. A paradigmatic example was the flyPAD, a new method for quantifying feeding behaviour in Drosophila melanogaster by measuring changes in electrode capacitance induced by the proboscis extension of a fly^12^. The integration of the flyPAD in Bonsai allowed researchers to quickly get started using this approach to setup new experiments. Furthermore, it also allowed the validation of the tool by enabling simultaneous acquisition of high-speed video recordings of fly behaviour which were later used for annotation and classification of the sensor feeding traces (Fig. 3g).

In a different set of experiments, Bonsai was used to implement a variation on a popular two-alternative forced choice (2AFC) decision-making task for rodents (Fig. 3h). In this family of tasks, animals are placed in an environment with three “ports”. They are presented with a stimulus in the centre port and afterwards report their perception of the stimulus by going either to the left or right choice ports. In the variation we present in this work, the two choice ports were replaced by regions of interest where the activity of the animal is analysed using computer vision. This example offered unique challenges as it combined sophisticated sequential control of a task environment with continuous data stream processing of video and sensor data.

The integration of all these diverse components for data acquisition and experiment control does not only allow for the rapid deployment of established protocols. In fact, the modular nature of their integration (i.e. how they can be combined together) opens up new avenues for research, by allowing a rich, rapid exploration of novel methodologies. To demonstrate this, we created a dynamic virtual environment for freely moving rodents where the visual presentation of a stimulus is tightly controlled in closed-loop to the actions of the animal. We used a projection setup similar to the low-cost multi-touch sensing table proposed by Han^13^, where a visible light rear-projection system is coupled with infrared illumination and an infrared imaging sensor to detect in real-time where the animal is located with respect to the visual display surface (Supp. Movie 1).

### Discussion

After about a year of using Bonsai in an active neuroscience research institute, dozens of different experimental protocols and data analysis pipelines have been successfully implemented using a subset of the provided building blocks^9^,^12^,^14^. We were surprised by the diversity of applications and by the pace at which new modules and devices were developed and integrated.

The performance achieved by Bonsai dataflow processing was an important consideration throughout. Video processing can be particularly challenging to handle given the bandwidth required to quickly acquire and process large data matrices. In order to correlate continuous measures of behaviour with neural activity, it is useful for those measurements to have both high spatial and high temporal resolution. Using Bonsai we were able to simultaneously process and compress grayscale image sequences from high resolution (1280×960) and high frame rate (120 Hz) cameras using standard off-the-shelf desktop computers (Intel Core i7, 8 GB RAM). In fact, many of the reported assays use multiple (>2) such video streams with success and actually process the behaviour video online either to control states of the behaviour protocol or to pre-process video data for offline analysis.

One of the areas where we see the application of Bonsai becoming most significant is in the development of dynamic behaviour assays (environments) using reactive control strategies. Brains evolved to generate and control behaviours that can deal with the complexity of the natural world. Unfortunately, when neuroscientists try to investigate these behaviours in the lab, they are often constrained by available technology in their efforts to reproduce equivalent environmental complexity in a controlled manner. As an example, consider a simple foraging scenario in which a land animal must collect, in a timely manner, food items that become available at random intervals in many sites. If the item is not collected in time, it rots or gets eaten by competitors. In the case of a single foraging site, a finite state machine description intuitively represents the workings of the environment (Fig. 4a). Let us now consider a situation where the environment has two of these food sites operating independently. Fig. 4b shows one possible, non-exhaustive model of such an environment. In the classical state machine formalism the machine can only be in one state at a time, which means that we now need to model this state as the combination of the individual states at each reward location. Furthermore, because transitions between these states are asynchronous and independent, we get edges in between nearly every pair of nodes, as each reward site can change its state at any point in time relative to the other.

**Figure 4:**
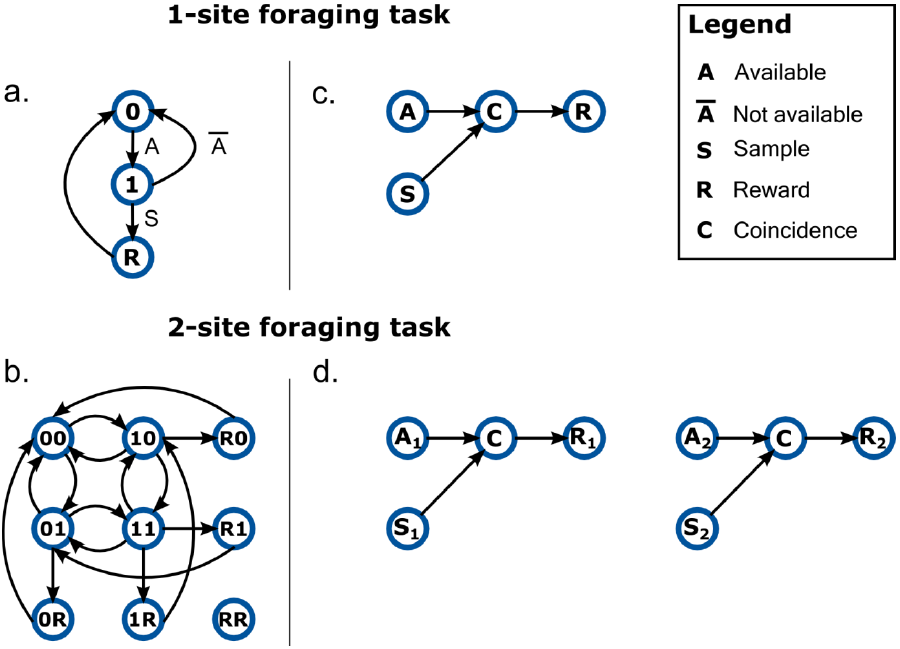
Describing the behaviour of dynamic environments using either state-machines or dataflows. **a.** A state-machine model of the 1-site foraging task. Zero indicates non-availability of reward at the site. One indicates reward is now available at the site. Labels on edges indicate event transitions. **b.** A non-exhaustive state-machine model for a foraging task with two sites. The active state is now a combination of the state of the two sites (indicated by a two character label) and all possible state combinations are tiled across the model. Event labels are omitted for clarity. Notation is otherwise kept. **c.** A dataflow model of the 1-site foraging task. Events in the state-machine model are now modelled as data sources. The coincidence detector node (C) propagates a signal only when the sample event closely follows reward availability. **d.** A dataflow model for a foraging task with two sites. The number subscripts denote foraging site index.

In order to make the design space more intuitive we need to introduce the parallel and asynchronous nature of the real-world situation into our modelling formalism. One simple example of this idea for the 1-site foraging task is depicted in Fig. 4c. In this case, we have two sources of events from the environment: one timer signalling the availability of reward (A); and a sampling event (S) which happens every time the animal checks the location for food. Both of these events can occur pretty much independently of each other, but when a sampling event coincides with reward availability (C), then reward (R) is delivered. Because this description is intrinsically asynchronous, it makes it extremely easy to scale the task to a larger set of locations: just replicate the dataflow for each of the other locations (Fig. 4d).

Another difficulty of the classical state machine formalism is dealing with continuous variables. The natural environment provides constant real-time feedback that tightly correlates with the actions of an animal. Reproducing such closed-loop interactions and manipulating their dynamics is a necessary tool for fully investigating brain function. Such models are virtually impossible to represent in a machine of finite states, given the potential infinitude of the feedback responses. However, the dataflow formalism of asynchronous event sources can easily accommodate such models. In fact, this is their natural battleground, where nodes represent reactive operators that promptly respond to input values broadcasted by event sources. These models of asynchronous computation are ideally suited to subsume the complexities of both discrete and continuous aspects of the natural environments that brains evolved to deal with, and will thus be required for neuroscientists to understand how the brain does so.

## Acknowledgments

We thank João Bártolo Gomes for suggesting the name Bonsai; Danbee Kim for early discussions on creating virtual environments for rodents; Joana Nogueira, George Dimitriadis and all the members of the Intelligent Systems Laboratory for helpful discussions and comments on the manuscript. We also thank all the members of the Champalimaud Neuroscience Programme who used Bonsai to setup their data analysis and acquisition experiments and in so doing provided valuable feedback to improve the framework. This project was supported by the NeuroSeeker Project Grant (FP7-ICT-2011-9:600925) and the Bial Foundation. GL is supported by the PhD Studentship SFRH/BD/51714/2011 from the Foundation for Science and Technology. The Champalimaud Neuroscience Programme is supported by the Champalimaud Foundation.

### Author Contributions

Conceived and developed the project: GL, NB, JF; Developed software: GL; Conceived and developed the experiments: GL, NB, JF, JPN, BVA, SS, LM, SM, PMI, PAC, REM, LC, ED, JJP, ARK; Performed and analysed experiments: GL, JF, JPN, BVA, SS, LM, SM, PMI, PAC, REM, LC, ED; wrote the manuscript: GL, ARK.

### Competing Financial Interests Statement

The authors declare no competing financial interests.

## Materials and Methods

The Bonsai framework can be freely downloaded from https://bitbucket.org/horizongir/bonsai.

All experiments were approved by the Champalimaud Foundation Bioethics Committee and the Portuguese National Authority for Animal Health, Direc?ao-Geral de Alimenta?ao e Veterinaria (DGAV).

### High-speed tracking of zebrafish behaviour

Larval zebrafish (6 dpf) were filmed with a high-speed monochrome video camera (Flea3, Point Grey, CA) under IR illumination. Fish swam freely in a custom-built arena that was laser cut from transparent acrylic that consisted of 3 separate chambers, each 40 × 100 mm. The position and orientation of the zebrafish in the central chamber was continuously tracked in real-time, while the video of the entire arena (1.3 Megapixel) was compressed to a high-quality H.264 encoded file that was used for subsequent offline analysis of the behaviour of a group of zebrafish placed in either of the side chambers.

### Mouse tracking and bulk fluorescence measurement of neuronal calcium activity

Freely behaving mice were filmed with a video camera (PlayStation Eye, Sony, JP) under white light illumination in their own homecages. A fiberoptic setup was developed to monitor bulk fluorescence changes in specific neuron populations using genetically encoded calcium indicators. Changes in fluorescence caused by neuronal activity were transmitted by an optical fiber and recorded with a CCD camera (Pike, Allied Vision Technologies, DE). The position and orientation of the mice was continuously tracked in real-time, while the mean pixel value of the area of the camera facing the fiber optic was continuously calculated. Both videos were compressed to high-quality H.264 encoded files to be used in subsequent offline analysis.

### Tracking human behaviour during a stochastic sound discrimination task

Bonsai was used to acquire timestamped images from 3 cameras (eye, person’s view, and arm) simultaneously. The videos are synced with presented sound stimulus by using the arm camera to also capture 2 IR LEDs which are turned on at sound on-set and off-set, respectively. The arm is tracked by using an IR LED mounted on a joystick and processing the video online at 120 Hz to minimize noise from compression. All the videos are compressed to a MP4 encoded file for offline analysis of the eye movements, pupil dilation, and syncing of all events with the sounds. The eye videos are captured at 30 Hz using the IR illuminated pupil headset https://code.google.com/p/pupil/.

### 3D tracking of rodent head position

Adult mice performing a two alternative forced choice task were filmed with a high-speed monochrome video camera (Flea3, Point Grey, CA). A fiber optic cable was attached to the mouse’s head. The 3D position and orientation of the head was tracked in real-time using square fiducial markers from the ArUco tracking library (Munoz-Salinas, 2014). The video (0.24 Megapixel) was simultaneously compressed to a high-quality H.264 encoded file that was used for subsequent offline analysis of the behaviour.

### Real-time stimulation conditioned to a region in space

Black mice were recorded with a high speed video camera (Flea3, Point Grey, CA), while exploring an open field arena (50×40 cm, L×W), under white illumination (~250 lux). The x and y position, body centroid and orientation of the animal in the arena was continuously tracked in real-time. Mice were implanted with an optical fiber connected to a laser, in order to receive photostimulation with blue light. A region of interest (ROI, 13×10.5 cm, L×W) was defined and a python script was written to outline the conditioning protocol. A digital output signal was sent to a microcontroller board (Uno, Arduino, IT), each time the body centroid of the animal entered in the ROI, producing photostimulation. All data for the animal tracking and digital output was saved in a .csv file, as well as the video, for subsequent offline analysis of the behaviour.

### Acute recordings from dense silicon probes

Recordings of spontaneous neural activity in motor cortex were performed in anesthetized rodents by means of silicon probes comprising a dense electrode array (A1×32–Poly35mm–25s– 177–CM32, Neuronexus, US). An open-source electrophysiology acquisition board (Open Ephys) was used along with a RHD2000 series digital electrophysiology interface chip that filters, amplifies, and digitally multiplexes 32 electrode channels (Intan Technologies, US). Extracellular signals sampled at 30 kHz with 16–bit resolution in a frequency band from 0.1 to 7500 Hz were saved into a raw binary format for subsequent offline analysis. Online analysis of neural spike waveforms across all probe channels was performed by aligning the multi-channel raw data on spike events from a selected channel of interest. A custom Bonsai visualizer was written using OpenGL to display all channel traces superimposed on the geometric arrangement of probe sites. It was possible to examine the details of extracellular activity in the spatial distribution.

### Recording drosophila feeding behaviour

Individual Drosophila melanogaster flies were allowed to freely feed on the flyPAD (Itskov, 2014) while their feeding behaviour was monitored at 50 Hz with a video camera (Flea3, Point Grey, CA) mounted on a Zeiss Discovery v.12 Stereo Microscope (Carl Zeiss, DE). FlyPAD measures fly’s behaviour on the food source by recording the capacitance at 100 Hz between the electrode on which the fly stands and the food. Videos were compressed to high-quality H.264 encoded files and subsequently manually annotated by a human observer to be used as a benchmark for the development of the automatic algorithms for the extraction of feeding behaviour from the capacitive trace. For more info, visit http://flypad.pt/.

### 2AFC task using video triggered reward

Adult PWD female mice were tested in behavioural experiments using restricted social interaction with adult C57BL6 and PWK males as reward. The behavioural paradigm consists of a custom built arena made of modular acrylic pieces assembled in an aluminum frame. The contact zone between the female and the male (composed of four holes with r = 0.5 cm) was either available for a fixed period of time, or physically restricted by a vertically sliding door controlled by a servomotor. Subjects initiated the interaction by nose-poking in an infrared beam port that would trigger the opening of the door and subsequent availability of the contact zone. Videos were recorded using high-speed monochrome video cameras (Flea3, Point Grey, CA). Performance, monitoring and control of the behaviour box was done using a Motoruino board (Motoruino, Artica, PT) and custom Bonsai scripts.

### Dynamic virtual environment for freely moving rodents

Three to five months old Long-Evans female rats were trained sequentially to forage and hunt virtual elements in a projected display, in exchange for water rewards. The behavioural paradigm consists of a custom built arena made of structural framing components (Bosch Rexroth, DE). The floor of the arena is a rear-projection screen made out of a frosted acrylic panel. In order to compensate for the short-distance, the projected image is reflected off a mirror positioned below the arena floor. Video was recorded using a high-speed monochrome video camera (Flea3, Point Grey, CA) equipped with a visible light cutoff filter (R72, Hoya, JP) and analysed in real-time using Bonsai. Infrared LED strips were positioned at the bottom of the arena in order to illuminate the floor through the diffuser, allowing for the tracking of the animal without contamination from the visual stimulus. Animals were first conditioned to a tone as a secondary reinforcer and then subsequently trained to either touch the light presented at random locations (foraging) or pursue a moving spot (hunting). Performance, monitoring and control of the behaviour box was done using an Arduino board (Micro, Arduino, IT) and a Bonsai reactive state machine.

